## Supplementary Information for "GPI-anchored FGF directs cytoneme-mediated bidirectional signaling to self-regulate tissue-specific dispersion"

**<u>Contents</u>**

**1. Supplementary Figures with Figure Legends (Suppl. Fig.1-Suppl. Fig.7)**

**2. Supplementary Tables 1-4.**

**3. Supplementary Notes:**

- **A. Expression analyses of *bnl* splice variants**
- **B. Bioinformatic analyses of physico-chemical properties of various constructs**
- **C. Comparison of *bnl-gal4*-driven expression of transgenic constructs**
- **D. Examples of gating strategy for FACS analyses**

### Supplementary Figure 1

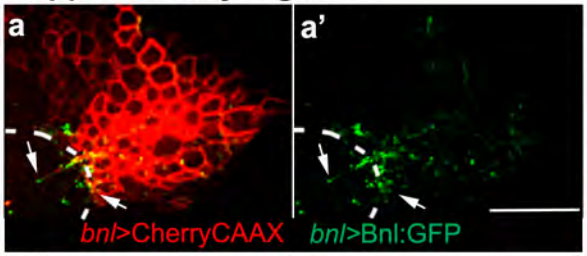


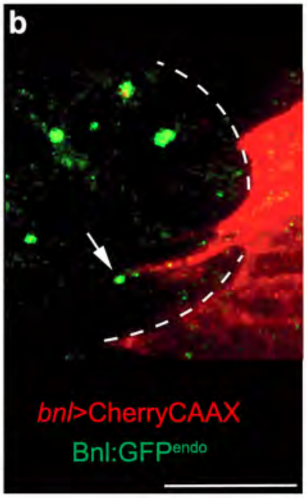


**c**

| | Sample # | Av. # cytonemes | |
|---|---|---|---|
| | | ASP (*btl-Gal4; UAS-CD8:GFP*) | Disc *bnl* source (*bnl-LexA; lexO-CherryCAAX*) |
| Live tissue | 5 | 31.6±1.7 | 7.6±2.5 |
| Fixed tissue | 5 | 11.2±2.8 | 2±1 |

**d**

| # of clones | # of cells in clones | # of cytonemes | # of cytonemes/cell |
|---|---|---|---|
| 1 | 1 | 2 | 2 |
| 2 | 1 | 2 | 2 |
| 3 | 5 | 19 | 3.8 |
| 4 | 2 | 7 | 3.5 |
| 5 | 1 | 5 | 5 |
| 6 | 1 | 5 | 5 |
| 7 | 2 | 6 | 3 |
| 8 | 8 | 20 | 2.5 |
| 9 | 2 | 16 | 8 |
| 10 | 2 | 2 | 1 |
| 11 | 2 | 7 | 3.5 |
| 12 | 2 | 5 | 2.5 |
| 13 | 1 | 8 | 8 |
| Average | 2.3 | 8 | 3.8 |

*btl*>CD8:GFP *bnl*>CherryCAAX

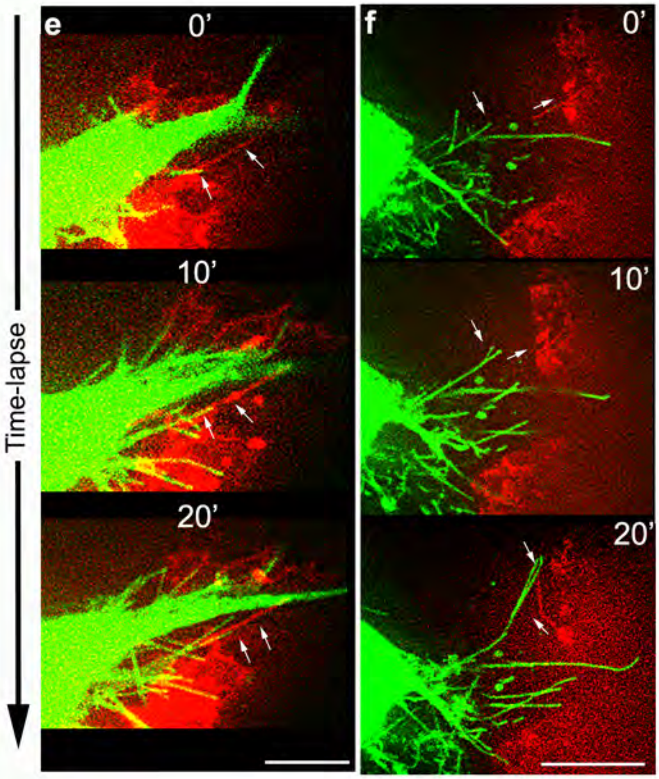


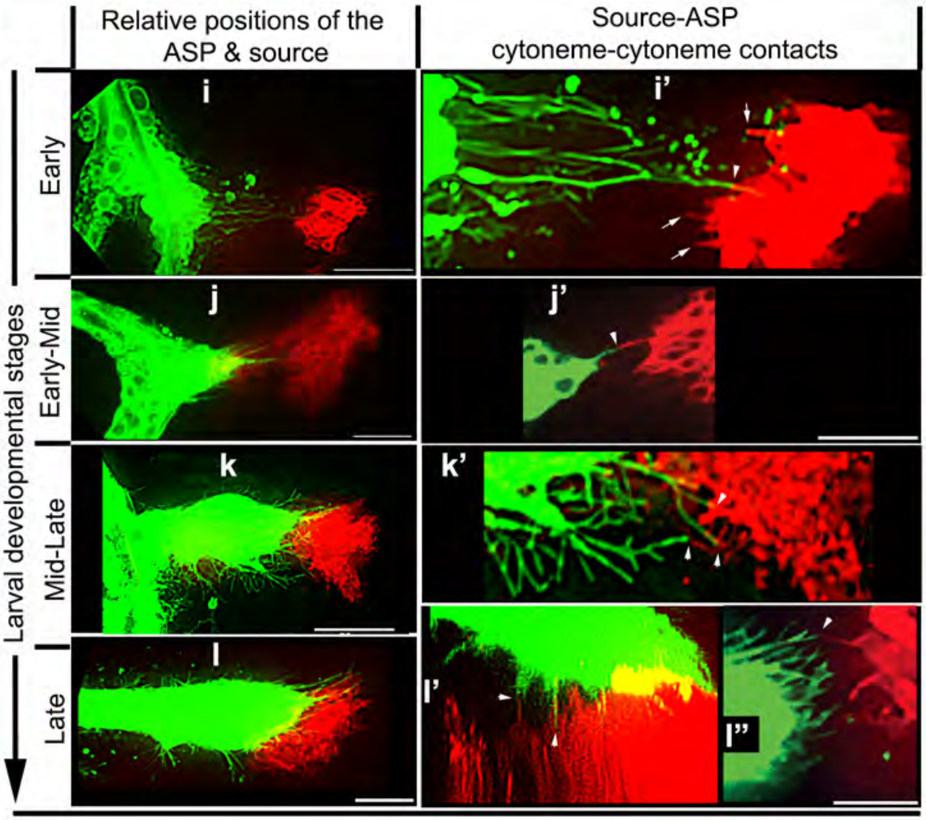


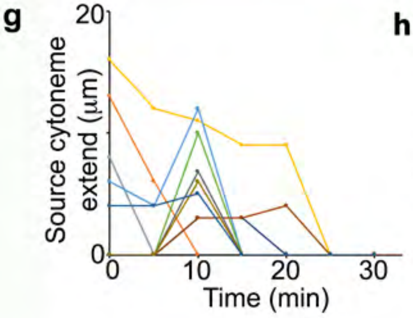


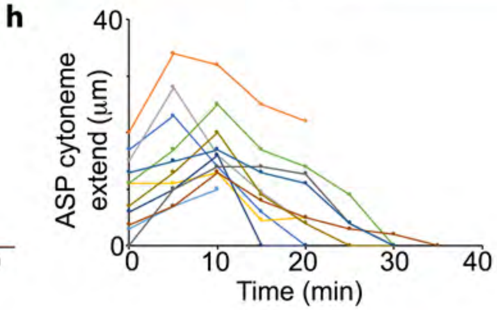

**Supplementary Figure 1. Bnl sending and receiving cytonemes reciprocally guide each other.**
**a-b** Live images of mCherryCAAX-marked wing disc *bnl* source (red) expressing either Bnl:GFP by *bnl-Gal4* (a,a'; *UAS-mCherryCAAX; bnl-Gal4* X *UAS-Bnl:GFP*) or endogenous Bnl:GFP$^{endo}$ (b; *bnl:gfp*$^{endo}$*/bnl>mCherryCAAX*; spinning disc confocal), showing ASP (dashed line)-specific polarity of Bnl:GFP presentation through cytonemes (arrows). **c** A comparison of cytoneme numbers (average ± S.D.) from the ASP and wing disc source under live and fixed imaging conditions, showing that the source cytonemes are detected mostly in live imaging. **d** Table showing the number of source cytonemes emanating from CD8:GFP-marked clones within the *bnl*-source (see Methods and Fig. 1h-h''). **e-h** Time-lapse images, showing repeated cycles of extension and retraction of source (red) and recipient (green) cytonemes for reciprocal contacts; g,h, Line plots showing interacting source and recipient cytoneme dynamics (also see Supplementary Table 1); the same color in g and h represents a pair of interacting source and ASP cytonemes. **i-l''** Maintenance of a convergently polarized cytoneme-forming niche at the ASP:source interface throughout the larval development. **e-l''** genotype - *btl-Gal4,UAS-CD8:GFP/+*; *bnl-LexA,lexO-mCherryCAAX/+*. Scale bars, 20 μm. Source data are provided as a Source Data file.

#### Supplementary Figure 2

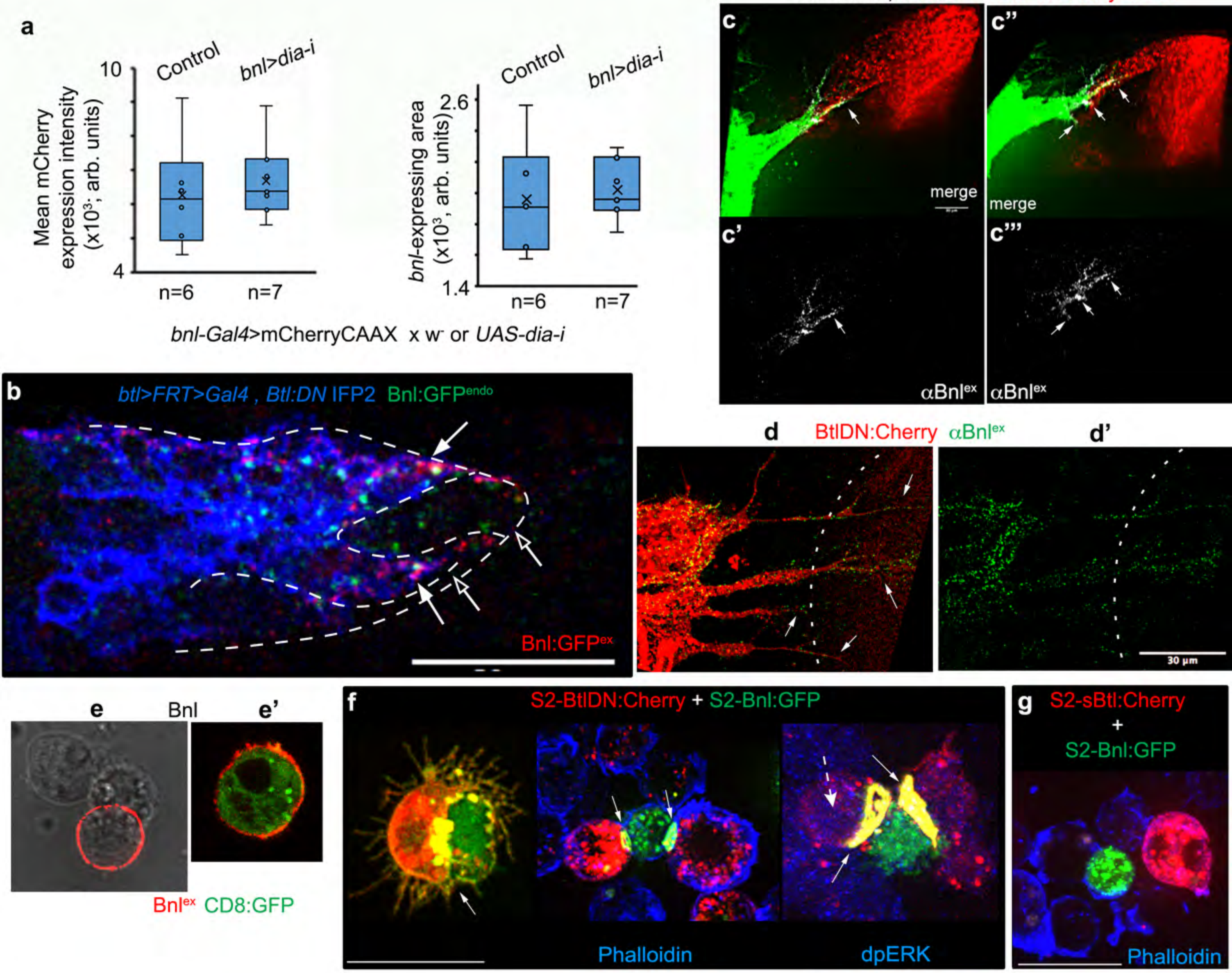


**h**

| S2 cells co-incubation | % trans-paired cells Btl-Bnl | Btl-Btl | Bnl-Bnl | # transfected cells |
|---|---|---|---|---|
| Btl:Cherry + Bnl:GFP | 30 | 4 | 2 | 1368 |
| BtlDN:Cherry + Bnl:GFP | 40 | 5 | 0 | 1026 |
| sBtl:Cherry + Bnl:GFP | 0 | 0 | 6 | 892 |

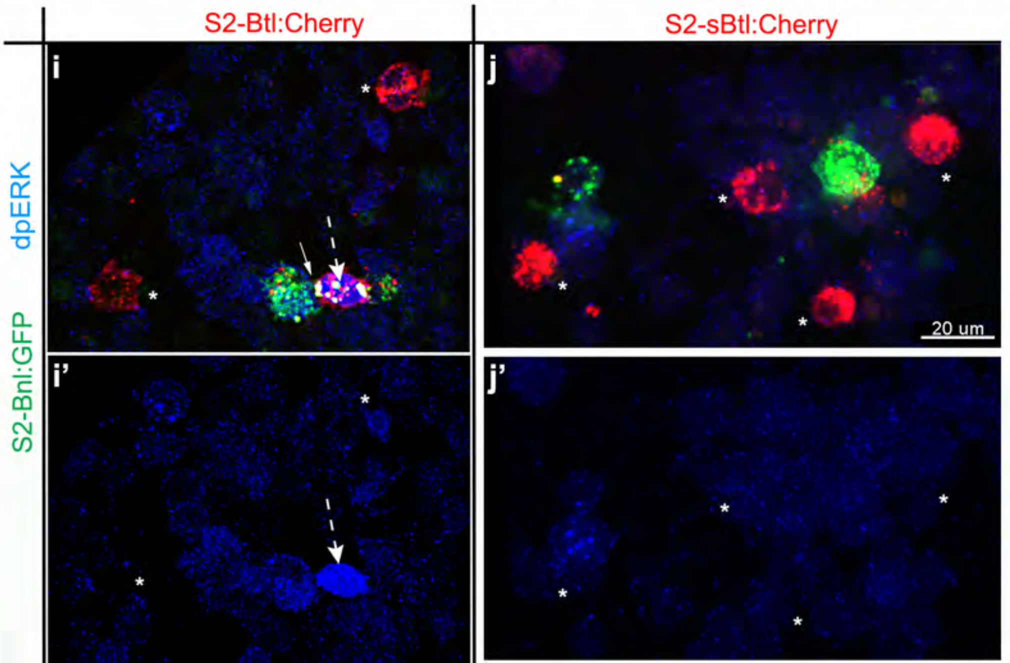

**Supplementary Figure 2. CAM-like Btl-Bnl binding mediates reciprocal contact formation.**
**a** *dia-i* expression under the *bnl-Gal4* control did not change *bnl* expression area or levels as detected by the mCherryCAAX expression (*bnl-Ga4, UAS-mCherryCAAX x UAS-dia-i* (for control, *w-* was crossed to *bnl-Ga4, UAS-mCherryCAAX*); in box plots, box shows the median as well as $1^{st}$ quartile and $3^{rd}$ quartile, and whiskers are minimum and maximum; n - biologically independent sample number; p values - unpaired two-tailed t-test; p=0.59 for mean mCherry expression intensity and p=0.72 for *bnl*-expressing area. **b** A mosaic ASP with Btl:DN-expressing IFP2-marked clones in the *bnl:GFP$^{endo}$* knock-in background (see Methods); Btl:DN-expressing surface areas on the ASP showed increased Bnl:GFP$^{endo}$ reception from the wing disc (arrow; probed by $\alpha$GFP$^{ex}$) compared to the WT areas (unmarked) of the same ASP tip (open arrow). **c-c'''** Bnl$^{ex}$ (grey, $\alpha$Bnl$^{ex}$) is asymmetrically enriched (arrow) at the contact sites between source and Btl:DN-expressing ASP projections or cytonemes; c''/c''', 3D projection of c/c'. **d,d'** Btl-DN:Cherry-containing cytonemes (arrow) emanating from a rudimentary ASP localized Bnl$^{ex}$ (green, $\alpha$Bnl$^{ex}$) puncta on their surfaces; dashed line, source area. **e,e'** $\alpha$Bnl$^{ex}$-stained S2 cells expressing either Bnl (e) or Bnl and CD8:GFP (e'; *act-Gal4*, *UAS*-Bnl, *UAS*-CD8:GFP), showing surface localized Bnl$^{ex}$, exclusively on the producing cell. **f** Different trans-paired forms of S2-Bnl:GFP and S2-Btl:DN:Cherry; arrow, trans-synaptic receptor-ligand co-clusters; dashed arrow, absence of nuclear dpERK in trans-paired S2-Btl:DN:Cherry. **g** Absence of trans-paring between S2-sBtl:Cherry and S2-Bnl:GFP. **h** Relatively high frequency of heterotypic Btl-Bnl trans-pairing in comparison to homotypic Btl-Btl or Bnl-Bnl trans-pairing in S2-Btl:Cherry variants/S2-Bnl:GFP co-incubation assays. **i-j'** Representative examples of $\alpha$dpERK-stained (blue) image frames, comparing trans-pairing experiments between S2-Bnl:GFP/S2-Btl:Cherry (i,i') and S2-Bnl:GFP/S2-sBtl:Cherry (j,j'); arrow, trans-synaptic Btl-Bnl co-cluster; dashed arrow, nucleus in trans-adhered Btl-expressing cells; *, receptor-expressing cells lacking dpERK. Scale bars, 20 $\mu$m, 30 $\mu$m (b,d,d'). Source data are provided as a Source Data file.

#### Supplementary Figure 3

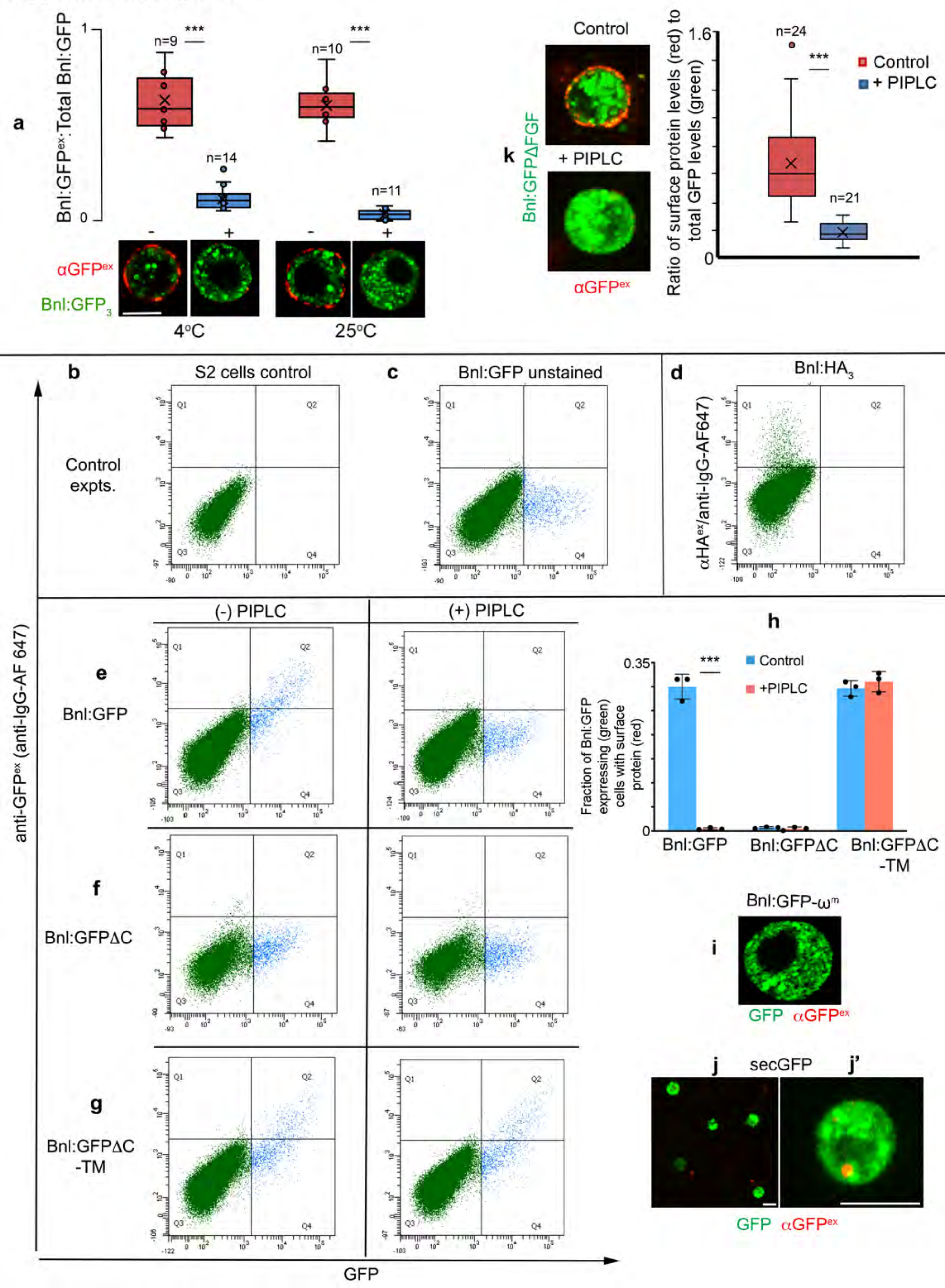

**Supplementary Figure 3. A GPI anchor tethers Bnl to the source cell surface.**

**a-h** S2 cells co-transfected with *actin-Gal4* and *UAS*-X; (X = Bnl:$HA_3$, Bnl:$GFP_3$, Bnl$HA_1$:$GFP_3$ (Bnl:GFP), Bnl:$HA_1$:$GFP_3\Delta$C (Bnl:GFP$\Delta$C), or Bnl:$HA_1$:$GFP_3\Delta$C-TM (Bnl:GFP$\Delta$C-TM) as indicated; cells were surface immune-stained either with HA or GFP antibodies as indicated. a, Box plots depicting the ratio of surface localized Bnl:$GFP_3$ (red, $\alpha GFP^{ex}$ immunostaining) to total Bnl:$GFP_3$ (green) in S2 cells before (-) after (+) the PIPLC treatment at various temperatures; lower panels, representative images of S2 cells as indicated; box shows the median as well as $1^{st}$ quartile and $3^{rd}$ quartile, and whiskers are minimum and maximum; n represents # cells examined; p values were calculated using unpaired two-tailed t-test; ***, p<0.001. **b-g** Representative flow cytometry profiles of S2 cells (b, control) or S2 cells expressing various constructs as indicated; b,c,d, FACS control; d, surface $\alpha HA^{ex}$ staining for Bnl:$HA_3$ as a control profile for GFP-positive cells. **h** Bar graphs showing quantitative values obtained from flow cytometry experiments depicting the average fraction of different Bnl:GFP variant-expressing cells (GFP positive) that contained surface localized signal ($\alpha GFP^{ex}$ immunostained (red)) before (control) and after the PIPLC treatment; values represent the mean ± SD from 3 independent transfection repeats; total number of GFP+ events examined over 3 independent experiments: 3670 (Bnl:GFP, control), 3095 (Bnl:GFP, +PIPLC), 3240 (Bnl:GFP$\Delta$C, control), 3044 (Bnl:GFP$\Delta$C, +PIPLC), 3000 (Bnl:GFP$\Delta$C-TM, control), 3000 (Bnl:GFP$\Delta$C-TM, +PIPLC); ***, p<0.001; p values were calculated using unpaired two-tailed t-test. **i** Lack of surface-localized protein (red, probed with $\alpha GFP^{ex}$) of Bnl:GFP-$\omega^m$ expressed in S2 cells. **j,j'** $\alpha GFP^{ex}$ immunostained S2 cells expressing secGFP construct showing the lack of surface distribution of the proteins due to its immediate secretion [1]. This is a control for bGFP-GPI, which is the same secGFP with Bnl's C-terminal signal sequence, leading to its GPI-anchoring to the producing cell surface (see Figure 4d'-f). **k** $\alpha GFP^{ex}$ immunostained S2 cells (left panels) expressing Bnl:GFP$\Delta$FGF construct showing PIPLC-sensitive surface distribution of the protein; right panel, box plots comparing the fraction of expressed protein on cell surface, with and without PIPLC treatment; box shows the median as well as $1^{st}$ quartile and $3^{rd}$ quartile, and whiskers are minimum and maximum; n represents # cells examined; p values were calculated using unpaired two-tailed t-test; ***, p<0.001. Scale bars, 10 μm. Source data are provided as a Source Data file.

#### Supplementary Figure 4

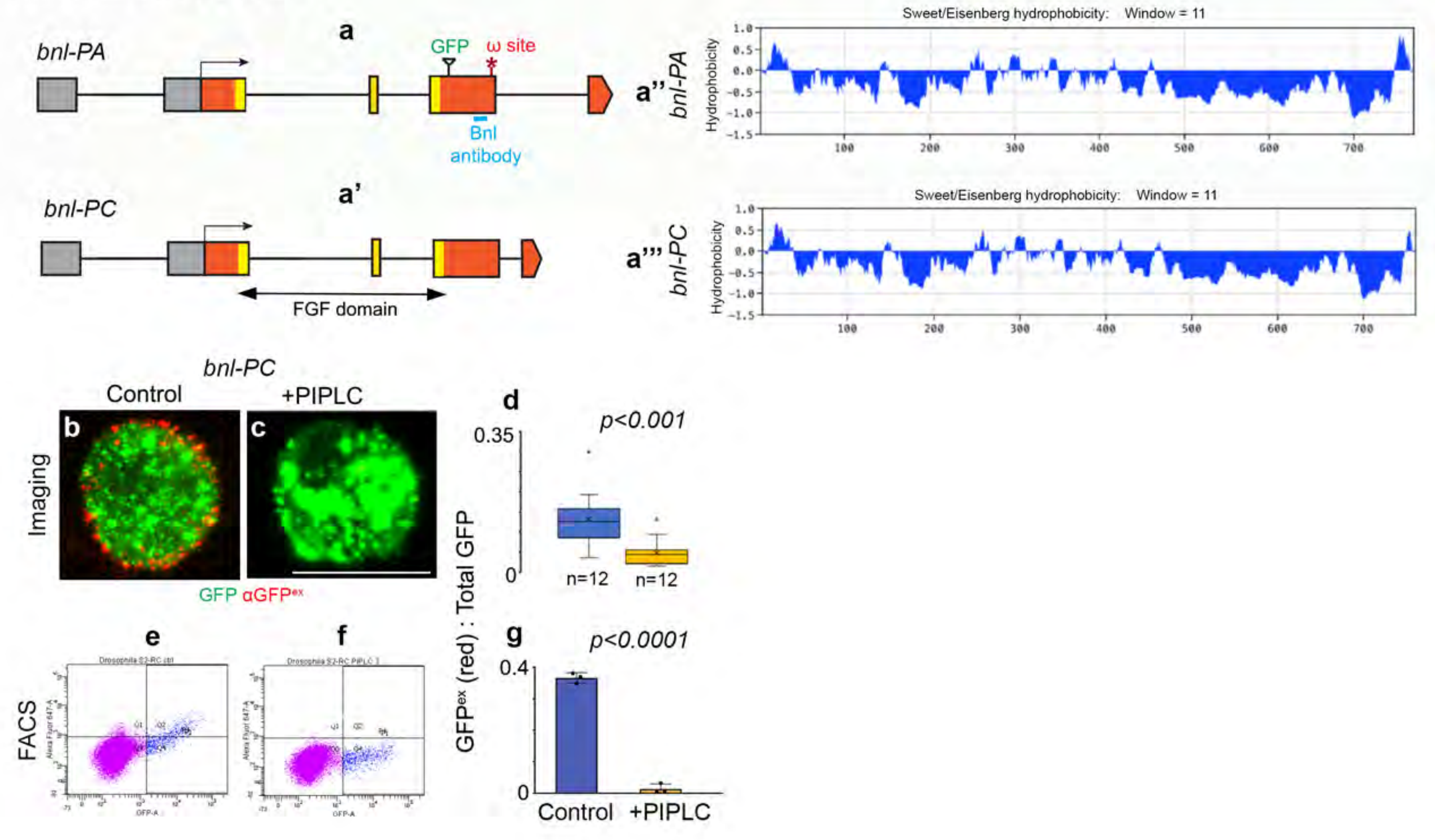

**Supplementary Figure 4. Characterization of GPI-anchored Bnl-PC isoform.**

**a-a'''** Comparison of PA and PC splice variants of *bnl*: **a,a'** Schematic maps of PA and PC loci highlighting identical FGF signaling domain, Bnl antibody binding site used for probing $Bnl^{ex}$, GFP tag (for probing $Bnl:GFP^{endo}$), putative $\omega$-site; Grey box, non-coding exons, colored box, coding exons. However, due to the alternative splicing of the last coding exons, *bnl-PC* isoform is 11 amino acid shorter at its C-terminal end than the *bnl-PA.* Subsequent to the common sequence between PA and PC, PC isoform has only 7 amino acids at its C-terminus, which is replaced by 18 amino acids sequences in PA. **a'',a'''** Comparative hydrophobicity plots of PA and PC variants showing a reduced hydrophobic stretch of C-terminal region of PC. **b-g** Extracellular $\alpha GFP^{ex}$-immunostaining of S2 cells expressing $Bnl:GFP_3$-PC showed surface-localized $Bnl:GFP_3$-$PC^{ex}$ (red), which is removed by PIPLC assay; **d** box plots showing surface localized fractions (red, probed with $\alpha GFP^{ex}$) of total $Bnl:GFP_3$-PC expressed in S2 cells before and after PIPLC treatment; box shows the median as well as 1$^{st}$ quartile and 3$^{rd}$ quartile, and whiskers are minimum and maximum; n represents # cells examined by imaging as shown in b,c; p value was calculated using unpaired two-tailed t-test; p=0.000649. **e,f** flow cytometric analyses of the same; g, bar graphs showing quantitative values obtained from flow cytometry experiments depicting the average fraction of $Bnl:GFP_3$-PC-expressing cells (GFP positive) containing surface localized $Bnl:GFP_3$-$PC^{ex}$ ($\alpha GFP^{ex}$ immunostained (red)) before and after the PIPLC treatment; values represent the mean $\pm$ SD from 3 independent experiments; total GFP+ events examined over 3 repeats: 3134 (control) and 2784 (+PIPLC); p value was calculated using unpaired two-tailed t-test. Scale bars, 10 $\mu$m. Source data are provided as a Source Data file.

### Supplementary Figure 5

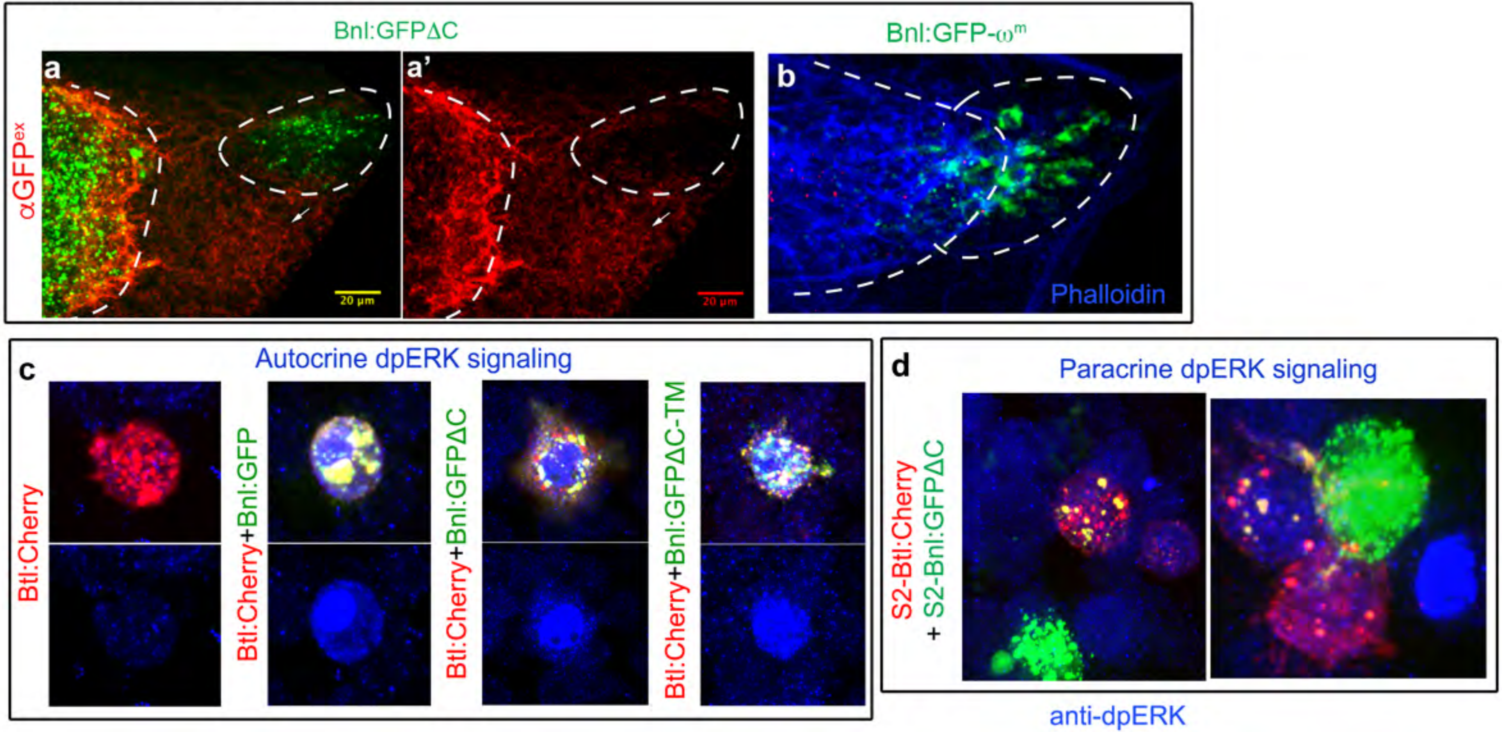

**Supplementary Figure 5. Autocrine and paracrine activity of Bnl variants.**
**a-b** Extracellular distribution of Bnl:GFP$\Delta$C (a,a') and Bnl:GFP-$\omega^m$ (b), when expressed under *bnl-Gal4* in the wing disc source; $\alpha$GFP$^{ex}$ immunostaining (red) showing that Bnl:GFP$\Delta$C is poorly retained on the source cell surface area (green punctate demarcated by dashed line), but are spread on the extracellular plane of the non-expressing disc cells (only red). ASP had both Bnl:GFP$\Delta$C$^{ex}$ and internalized Bnl:GFP$\Delta$C (probed only by GFP), showing non-autonomous signal dispersal. In contrast, Bnl:GFP-$\omega^m$ (b) is poorly externalized from the source and not received by the ASP (Phalloidin-stained). **c** Efficient autonomous MAPK signaling (nuclear dpERK, blue) of different Bnl:GFP variants when co-expressed with Btl:Cherry in S2 cells. **d** Inefficient non-autonomous MAPK signaling of Bnl:GFP$\Delta$C when S2-Bnl:GFP$\Delta$C cells were co-incubated with S2-Btl:Cherry cells.

#### Supplementary Figure 6

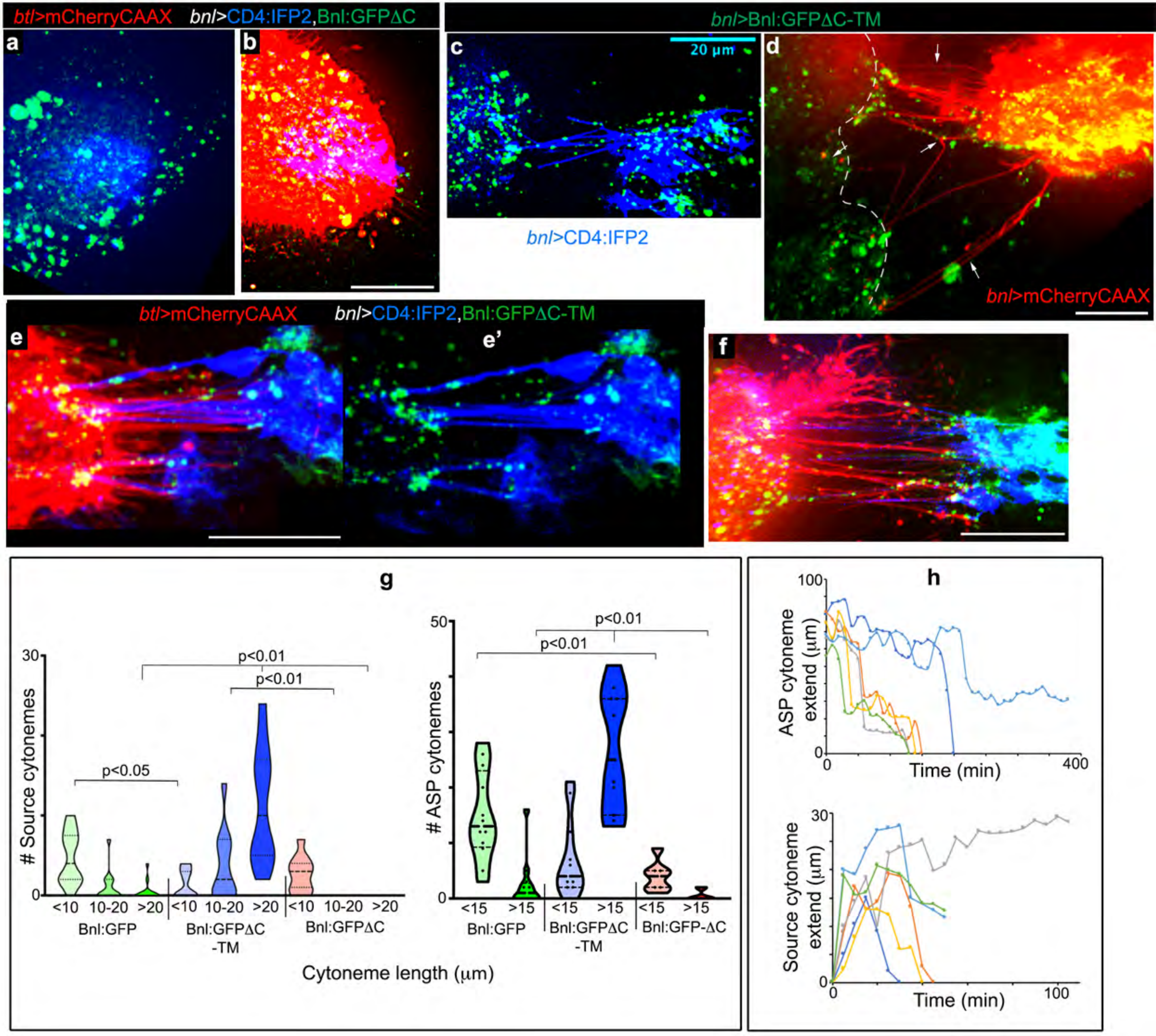

**Supplementary Figure 6. GPI-anchored Bnl induces cytoneme-mediated bidirectional matchmaking for contacts.**

**a** Split channels of Figure 6g, showing the random spread of Bnl:GFPΔC from its source (blue, *bnl>CD4:IFP2*). **b** An example showing the loss of ASP cytonemes (red), when Bnl:GFPΔC was overexpressed from CD4:IFP2-marked source cells (blue). **c** Split green and blue channels of Figure 6j showing Bnl:GFPΔC-TM-containing source cytonemes contacting the ASP. **d-f** Examples showing CAM-like activity of Bnl:GFPΔC-TM when expressed from the source: (d) Long polarized Bnl:GFPΔC-TM-expressing source cytonemes (arrows) (red, *bnl>mCherryCAAX*) connected to the ASP and disc-associated transverse connective (dashed outline). (e-f) Bundles of ASP and source cytonemes interacting through Bnl:GFPΔC-TM-enriched lateral contact sites; e', split blue and green channels of (e). **g** Violin plots showing a comparison of the number and length distribution of ASP and source cytonemes (from Fig. 6f-l) induced by Bnl:GFP, Bnl:GFPΔC, or Bnl:GFPΔC-TM, when overexpressed from the disc source; in violin plots, black dotted lines show the median as well as $25^{th}$ and $75^{th}$ percentiles; n=13 (Bnl:GFP, source), 11 (TM, source), 11 (ΔC, source), 12 (Bnl:GFP, ASP), 11 (TM, ASP), 7 (ΔC, ASP) biologically independent samples; p values were calculated using one way-ANOVA followed by Tukey's honestly significant different test. **h** Line plots showing dynamics of the source and recipient cytonemes as indicated when Bnl:GFPΔC-TM was expressed in *bnl* source using *bnl-Gal4* (see Supplementary Table 1); the same color represents the interacting Bnl-receiving and -sending cytonemes from the same sample. All panels, live imaging. Scale bars, 20 μm. Source data are provided as a Source Data file.

### Supplementary Figure 7

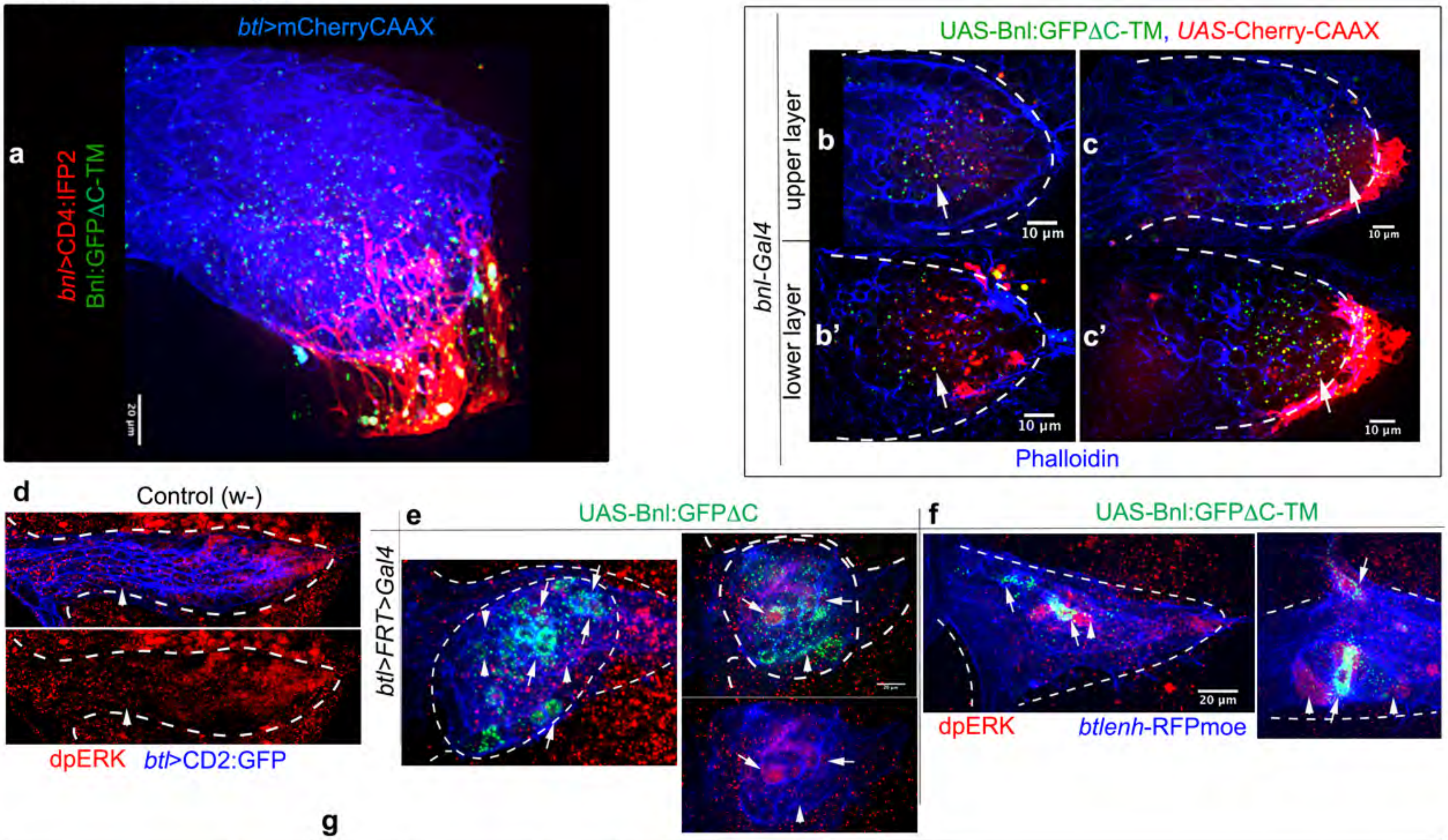


g

| Genotype *hsflp/+; btlenh>FRT>Gal4, btlenh-RFPmoe/UAS-"X"* | Sample # | Dispersal range (# cells) | % signal receiving ASP cells with nuclear dpERK | % clones induced growth | % clones induced branching |
|---|---|---|---|---|---|
| Bnl:GFP | 24 | 2.8±0.7 | 99.2±2.3 | 83 | 25 |
| Bnl:GFP-TM | 33 | 2.3±0.8 | 82.9±19.9 | 76 | 36 |
| Bnl:GFP-**Δ**GPI | 21 | 4.2±0.9 | 36.9±27.4 | 81 | 0 |

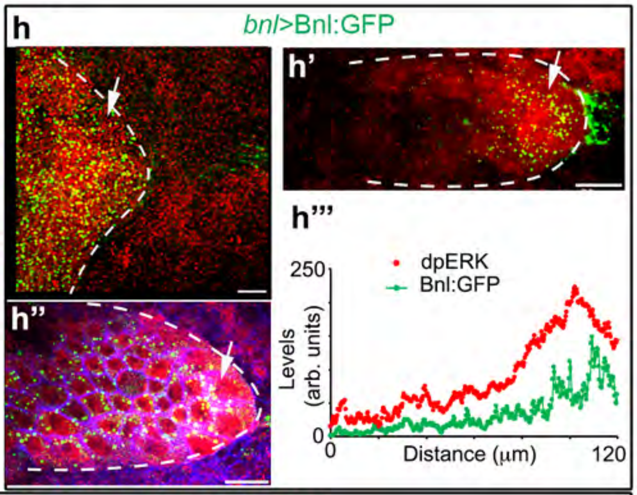


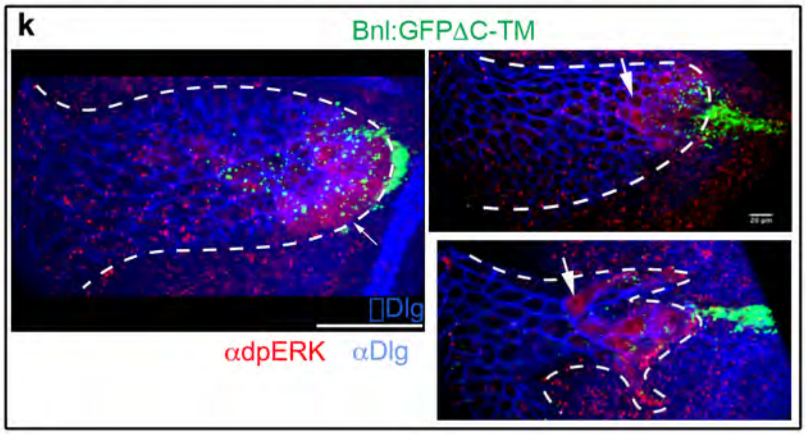


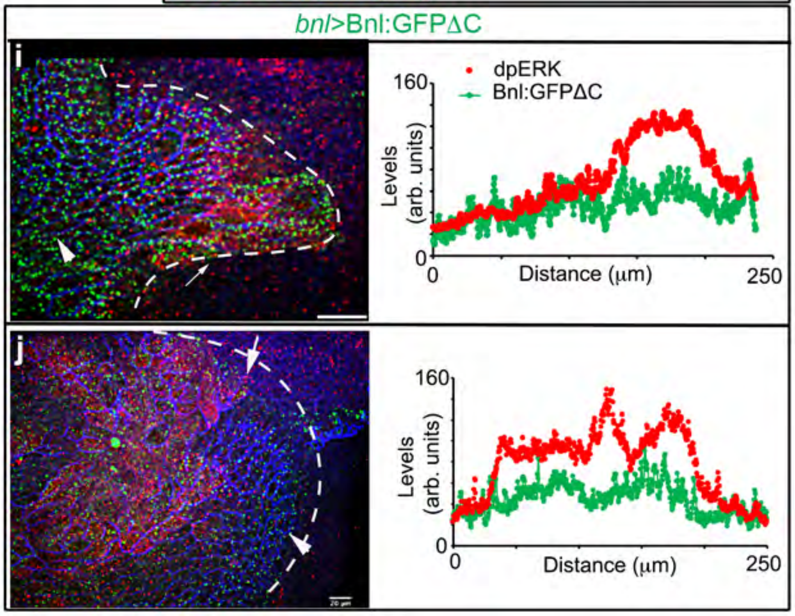


l

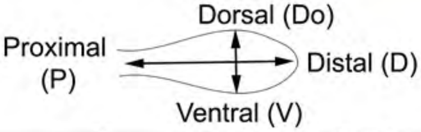


| Genotype | Sample # | Av. longest Do-V axis length | Av. longest D-P axis length | Ratio Do-V/D-P |
|---|---|---|---|---|
| *bnl:GFP$^{endo}$* * | 9 | 33±4 | 119±10 | 0.28±0.04 |
| *bnlGal4*>Bnl:GFP | 7 | 95±24 | 120±14 | 0.82±0.3 |
| *bnlGal4*>Bnl:GFP-TM | 15 | 56±11 | 137±10 | 0.41±0.07 |
| *bnlGal4*>Bnl:GFPΔGPI | 8 | 146±11 | 104±24 | 1.50±0.47 |

**Supplementary Figure 7. GPI anchoring is required for Bnl release and morphogen-like signaling.** **a** Strong affinity and adhesion of Bnl:GFPΔC-TM-expressing source cytonemes (red, *bnl>CD4:IFP2*) with the ASP surface (blue, *btl>mCherryCAAX*). **b-c'** Images of two wing discs expressing Bnl:GFPΔC-TM and mCherryCAAX from the *bnl* source (red), showing endocytosed Bnl:GFPΔC-TM puncta colocalized with the source membrane in upper and lower layer cells of the tubular ASP epithelium; Phalloidin-Alexa Fluor 647 (blue) marked cell outlines. **d** Control (*w-*) ASP showing the lack of nuclear dpERK in the ASP stalk and TC region, where the GOF clones were scored (see Fig. 9a-d). **e-g** Examples of ASPs with ΔC or TM GOF clones (1-2 cell size) (arrows) and their non-autonomous signaling (arrowhead; red, dpERK). **g** Table showing non-autonomous effects of Bnl:GFP, TM, and ΔC GOF clones in the ASP stalk; sample # (n) represents the number of clones examined over >10 biologically independent samples; values represent the mean ± SD; p values for % signal receiving ASP cells with nuclear dpERK: $p < 0.01$, for Bnl:GFP-ΔC vs. Bnl:GFP or Bnl:GFPΔC-TM; $p < 0.05$, for Bnl:GFP vs. Bnl:GFPΔC-TM; p values were calculated using one way-ANOVA followed by Tukey's honestly significant different test. **h-l** Comparative analyses of the activity of Bnl:GFP, Bnl:GFPΔC-TM (TM), and Bnl:GFPΔC (ΔC), expressed from wing disc source under *bnl-Gal4* (*bnl-Gal4 X UAS-X*)**:** (**h-h'''**) Wing discs expressing Bnl:GFP, showing a spatial coordination of signal distribution (green puncta), signaling patterns (dpERK, red), and ASP growth. (**i-k**) The coordination between signal distribution, signaling, and growth was uncoupled by ΔC expression and was regained with TM. However, TM distribution was restricted in range in comparison to Bnl:GFP. (**l**) Comparison of ASP shapes in conditions as indicated; *, homozygous *bnl:gfp^endo^* larvae used as the control for overexpressed Bnl:GFP variants; top panel, illustration showing the measurement of the longest Do-V and D-P axes (μm) from extended Z-projected ASP images; sample # (n) represents the number of biologically independent samples; values represent the mean ± SD; p values: Do-V/D-P axes: $p < 0.01$, *bnl:gfp*$^{endo}$ vs. Bnl:GFP or ΔC and Bnl:GFP vs. TM or ΔC; p values were calculated using one way-ANOVA followed by Tukey's honestly significant different test. h-k, arrows, recipient cells with signaling; arrowhead, recipient cells without signaling; αDlg (blue), cell outlines. All panels, dashed line shows ASP outline or ectopic tracheal outgrowth. Scale bars, 20 μm; 10 μm (b-c'). Source data are provided as a Source Data file.

#### Supplementary Tables

##### Supplementary Table 1. Quantification of the dynamics of the interacting cytonemes from Bnl-receiving and -sending cells.

| | FGF-receiving cells | | | FGF-sending cells | | |
|---|---|---|---|---|---|---|
| | WT | *bnl>Bnl:GFP* | *bnl>Bnl:GFP ΔC-TM* | WT | *bnl>Bnl:GFP* | *bnl>Bnl:GFP ΔC-TM* |
| Lifetime (min) | 19.55 ±6.1 | 17.5±5* | 180±101.39 | 7.73± 4.67 | ND | 46.67±30.11 |
| # Fluctuating peaks/lifetime | 1±0 | 1±0 | 4.33±2.66 | 1±0 | ND | 2.33±1.37 |
| Maximum extension (μm) | 19.36 ±7.4 | 15.8±3.8 | 76.67±8.98 | 8.35±4.04 | ND | 26.52±7.3 |
| Average extension rate (μm/min) | 1.28 ±0.79 | 0.85±0.32 | 0.47±0.15 | 1.13±0.57 | ND | 0.70±0.23 |
| Average retraction rate (μm/min) | 1.37 ±0.75 | 0.93±0.19 | 0.88±0.22 | 1.32±0.56 | ND | 0.79±0.26 |

Note: Maximum extension - the maximum length of a cytoneme during its lifetime. Peak - Each extension and retraction cycle within the lifetime of a cytoneme. Number of fluctuating peaks/lifetime - the number of extension and retraction cycles of a cytoneme. Average extension and retraction rates - measured by the net cytoneme length change/time during its extension or retraction.

*, For this condition, 4 long cytonemes were used. Most cytonemes (N=25) were short in length and had lifetime <10 min and was not counted in 10 min interval time-lapse movies.

Values represent mean ± SD. N=11 cytonemes for WT; 6 cytonemes for *bnl>Bnl:GFPΔC-TM* receiving and sending cytonemes; 4 cytonemes for *bnl>Bnl:GFP* receiving cytonemes. p values (WT vs *bnl>Bnl:GFPΔC-TM*) for receiving cytoneme dynamics: lifetime, $p < 0.0001$; # fluctuating peaks/lifetime, $p < 0.001$; maximum extension, $p < 0.0001$; average extension rate, $p = 0.025$; average retraction rate, no significant difference. p values for sending cytoneme dynamics: lifetime, $p < 0.001$; # fluctuating peaks/lifetime, $p = 0.0046$; maximum extension, $p < 0.001$; average extension rate, no significant difference; average retraction rate, $p = 0.047$. p values were calculated using unpaired two-tailed t test. $p < 0.05$ is considered significant. Source data are provided as a Source Data file.

<u>Genotypes</u>: WT: *btlGal4,UAS-CD8:GFP/+;bnlLexA,LexO-mCherryCAAX/+*.

*bnl>Bnl:GFP*: *btlLexA,LexO-mCherryCAAX/UAS-CD4:mIFP; bnlGal4/UAS-Bnl:GFP.*

*bnl>Bnl:GFPΔC-TM*: *UAS-Bnl:GFPΔC-TM/UAS-CD4:mIFP; bnlGal4/btlLexA,LexO-mCherryCAAX* for FGF-receiving cytonemes, and *UAS-mCherryCAAX/UAS-Bnl:GFPΔC-TM; bnlGal4/+* for FGF-sending cytonemes.

**Supplementary Table 2. Comparison of the ASP and source cytoneme numbers when *diaRNAi* was expressed in the ASP.**

| | | # ASP cytonemes < 15 μm | # ASP cytonemes > 15 μm | # FGF source cytonemes |
|---|---|---|---|---|
| Control (N=11) | Average | 11.55 | 17 | 9.45 |
| | SD | ±4.57 | ±4.96 | ±3.30 |
| *btlGal4>diaRNAi* (N=8) | Average | 10 | 1 | 0 |
| | SD | ±7.66 | ±1.06 | ±0.35 |
| *p* (unpaired t-test) | | 0.559367458 | 5.87525E-08 | 4.27454E-07 |

Note: Values represent mean ± SD. N represents the number of biologically independent samples. p values were calculated using unpaired two-tailed t test. Source data are provided as a Source Data file.
Genotypes: Control: *btlGal4,UAS-CD8GFP/+;bnlLexA,LexO-mCherryCAAX/+.*
*btl-Gal4>diaRNAi*: *btlGal4,UAS-CD8GFP/tub-Gal80*$^{ts}$*;bnl-LexA,LexO-mCherryCAAX/ UAS-diaRNAi.*

**Supplementary Table 3: Autocrine and paracrine MAPK signaling activity of Bnl:GFP variants in S2 cells**

**A. Autocrine activity: S2-Btl:Cherry co-expressed with Bnl:GFP variants**

| S2 cell co-expression | # total Btl cells | # Btl cells with nuclear dpERK | % Btl cells with nuclear dpERK |
|---|---|---|---|
| Bnl:GFP+Btl:Cherry | 15 | 14 | 93 |
| Bnl:GFPΔGPI+Btl:Cherry | 22 | 21 | 95 |
| Bnl:GFPΔGPI-TM +Btl:Cherry | 16 | 15 | 94 |

**B. Paracrine activity: S2-Bnl:GFP variants co-incubated with S2-Btl:Cherry**

| S2-Btl:Cherry cells (# or %) | Bnl:GFP +Btl:Cherry | Bnl:GFPΔGPI +Btl:Cherry | Bnl:GFPΔGPI-TM +Btl:Cherry |
|---|---|---|---|
| # of total cells (A) | 169 | 188 | 123 |
| # of (A) with dpERK | 76 | 65 | 60 |
| % of (A) with dpERK | 44.9 | 34.6* | 48.8 |
| # of trans-paired cells with S2-Bnl:GFP/TM or adjacent to S2-ΔC (B) | 78 | 33 | 87 |
| # of (B) with dpERK | 71 | 5 | 58 |
| % of (B) with dpERK | 91 | 15** | 67 |
| # of uncoupled Btl +ve cells (C) | 91 | 188 | 36 |
| # of (C) with dpERK | 5 | 60 | 2 |
| % of (C) with dpERK | 5 | 32 ** | 6 |

Note: data represents results from three independent transfection repeats.

#### Supplementary Table 4: Resources and reagents used in this study

| REAGENT or RESOURCE | DESCRIPTION | SOURCE |
|---|---|---|
| Antibodies | | |
| Mouse monoclonal anti-Discs large (Dlg) | 1:100 | DSHB, Cat# 4F3 anti-discs large; RRID: AB_528203 |
| Phospho-p44/42 MAPK (Erk1/2) (Thr202/Tyr204) rabbit monoclonal antibody (dpERK) | 1:250 in tissue and 1:1000 in S2 cells | Cell signaling Technology; Cat# 4370; RRID: AB_2315112 |
| Rat monoclonal anti-HA (3F10) | 1:1000 for standard and 1:500 for EIF | Roche; Cat#11867423001; RRID: AB_390918 |
| Rabbit polyclonal anti-Bnl | 1:500 for EIF [1] | N/A |
| Rabbit anti-GFP antibody | 1:3000 for EIF | Abcam; Cat# ab6556; RRID: AB_305564 |
| Goat anti-Mouse IgG (H+L), Alexa Fluor 555 | 1:1000 | Thermo Fisher Scientific; A21434 |
| Goat anti-Mouse IgG (H+L), Alexa Fluor 647 | 1:1000 | Thermo Fisher Scientific; A28181 |
| Goat anti-Rat IgG (H+L), Alexa Fluor 647 | 1:1000 | Thermo Fisher Scientific; A21247 |
| Goat anti-Rabbit IgG (H+L), Alexa Fluor 555 | 1:1000 | Thermo Fisher Scientific; A21428 |
| Goat anti-Rabbit IgG (H+L), Alexa Fluor 647 | 1:1000 | Thermo Fisher Scientific; A21244 |
| Bacterial and Virus Strains | | |
| DH5 Alpha | | |
| Chemicals, Peptides, and Recombinant Proteins | | |
| Alexa Fluor 647 Phalloidin | Thermo Fisher Scientific | Cat# A22287, RRID: AB_2620155 |
| Furin Inhibitor I - Calbiochem | Sigma-Aldrich | Cat# 344930 |
| Furin Inhibitor II - Calbiochem | Sigma-Aldrich | Cat# 344931 |
| Phospholipase C, Phosphatidylinositol-specific from *Bacillus cereus* | Invitrogen | Cat# P6466 |
| Critical Commercial Assays | | |
| Lipofectamine 3000 Transfection Reagent | Thermo Fisher Scientific | Cat# L3000008 |
| Mirus TransIT®-Insect Transfection Reagent | Mirus Bio | |
| TRI Reagent | Sigma-Aldrich | Cat# T9424 |
| One*Taq*® One-Step RT-PCR Kit | NEB | Cat# E5315S |
| Deposited Data | | |
| Raw data from all the figures | This paper | |
| Experimental Models: Cell Lines | | |
| *D. melanogaster*: Cell line S2: S2-DRSC | Laboratory of Thomas B. Kornberg | FlyBase: FBtc0000181 |
| Experimental Models: Organisms/Strains | | |
| *D. melanogaster*: *UAS-Bnl:GFPΔC* | This paper | N/A |
| *D. melanogaster*: *UAS-Bnl:GFPΔC-TM* | This paper | N/A |

| | | |
|---|---|---|
| *D. melanogaster*: *UAS-Bnl:GFPΔC*$_{168}$*-TM* | This paper | N/A |
| *D. melanogaster*: *UAS-Bnl:GFPΔC*$_{168}$ | This paper | N/A |
| *D. melanogaster*: *LexO-BtlDN:Cherry* | This paper | N/A |
| *D. melanogaster*: *UAS-Bnl:GFP−ω*$^{m}$ | This paper | N/A |
| *D. melanogaster*: *bnl:gfp*$^{endo}$ | [1] | N/A |
| *D. melanogaster*: *btl:cherry*$^{endo}$ | [1] | N/A |
| *D. melanogaster*: *UAS-Bnl:GFP* | [2] | N/A |
| *D. melanogaster*: *UAS-CD8:GFP* | BDSC | 5137 |
| *D. melanogaster*: *UAS-nlsGFP* | BDSC | 4776 |
| *D. melanogaster*: *UAS-mCherryCAAX* | BDSC | 59021 |
| *D. melanogaster*: *UAS-CD4:mIFP* | BDSC | 64182 |
| *D. melanogaster*: *lexO-mCherryCAAX* | [3] | N/A |
| *D. melanogaster*: *UAS-Btl*$^{DN}$ | [4] | N/A |
| *D. melanogaster*: *UAS-Bnl* | BDSC | 64232 |
| *D. melanogaster*: *UAS-diaRNAi* | BDSC | 33424 |
| *D. melanogaster*: *UAS-Dia-GFP* | [3] | N/A |
| *D. melanogaster*: *UAS-ΔDAD-Dia-GFP* | [3] | N/A |
| *D. melanogaster*: *bnl-LexA* | [1] | N/A |
| *D. melanogaster*: *bnl-Gal4* | BDSC | 112825 |
| *D. melanogaster*: *btl-Gal4* | [5] | N/A |
| *D. melanogaster*: *btl-LHG* | [3] | N/A |
| *D. melanogaster*: *hs-FLP; btl>y+>Gal4, btl-mRFP1moe* | [6] | N/A |
| *D. melanogaster*: *hs-mFlp* | BDSC | N/A |
| *D. melanogaster*: *FlyBow FB2.0* | BDSC | N/A |
| *D. melanogaster*: *btl:GFP* fTRG | VDRC | 318302 |
| *D. melanogaster*: *hs-Flp* | BDSC | 6 |
| *D. melanogaster*: *tub-Gal80*$^{ts}$ | BDSC | 7108 |
| *D. melanogaster*: *act>CD2>Gal4* | BDSC | 4780 |
| *D. melanogaster*: *w*$^{1118}$ | BDSC | 3605 |
| Oligonucleotides | | |
| Primer for cloning *UAS-Bnl:GFPΔC*: GCCAAGCTTGCATGCCGGTACCTTAGTAGCTCGCATCTTCTAGGGATCC | This paper | N/A |
| Primer for cloning *UAS-Bnl:GFPΔC-TM:* CCCTAGAAGATGCGAGCTACGACTTCGCCTGTGATATTTACATCTGG | This paper | N/A |
| Primer for cloning *UAS-Bnl:GFPΔC-TM:* GATGTAAATATCACAGGCGAAGTCGTAGCTCGCATCTTCTAGGGATCC | This paper | N/A |
| Primer for cloning *UAS-Bnl:GFPΔC-TM:* GCCAAGCTTGCATGCCGGTACCTTAGTGGTAGCAGATGAGAGTGATGATC | This paper | N/A |
| Primer for cloning *UAS-Bnl:GFPΔC*$_{168}$: GCCAAGCTTGCATGCCATATATTCTAGATTACTTCTTCTTGCCTCCGTGCTG | This paper | N/A |
| Primer for cloning *UAS-Bnl:GFPΔC*$_{168}$*-TM*: CAGCACGGAGGCAAGAAGAAGgacttcgcctgtgatatttacatctgg | This paper | N/A |
| Primer for cloning *UAS-Bnl:GFPΔC*$_{168}$*-TM*: ccagatgtaaatatcacaggcgaagtcCTTCTTCTTGCCTCCGTGCTG | This paper | N/A |
| Primer for cloning *UAS-Bnl:HA*$_{1}$*GFP*$_{3}$*Cherry*$_{c}$: GTTTTGCTCCGAAAAAGAGCCATCCTGATGGTGAGCAAGGGCGAGGAG | This paper | N/A |
| Primer for cloning *UAS-Bnl:HA*$_{1}$*GFP*$_{3}$*Cherry*$_{c}$: GCTGCTGGTACCTTACTTGTACAGCTCGTCCATGCCG | This paper | N/A |

| | | |
|---|---|---|
| Primer for cloning *UAS-Bnl:GFP-*$\omega^m$*:* CGAGGCCCAAGGACGCCCCCCACCAGGCGGCGACGAT TCG | This paper | N/A |
| Primer for cloning *UAS-Bnl:GFP-*$\omega^m$*:* CGAATCGTCGCCGCCTGGTGGGGGGCGTCCTTGGGCC TCG | This paper | N/A |
| Primer for cloning *UAS-bGFP-GPI:* CAACAACTTGACAATGTCCAAGGGCGAGGAG | This paper | N/A |
| Primer for cloning *UAS-bGFP-GPI:* GCCCTTGGACATTGTCAAGTTGTTGTCCATGGCC | This paper | N/A |
| Primer for cloning *UAS-bGFP-GPI:* TGGATGAGCTGTACAAGACCGAGGGCGACGGTG | This paper | N/A |
| Primer for cloning *UAS-bGFP-GPI:* TCGGTCTTGTACAGCTCATCCATGCCC | This paper | N/A |
| Primer for cloning *UAS-Bnl:GFPΔFGF:* CCCTTGGACATGGACTGTGGCACCGTGG | This paper | N/A |
| Primer for cloning *UAS-Bnl:GFPΔFGF:* TGCCACAGTCCATGTCCAAGGGCGAGGAGC | This paper | N/A |
| Primer for cloning *UAS-Bnl:GFPΔFGF:* CACCGTCTTGTACAGCTCATCCATGCCC | This paper | N/A |
| Primer for cloning *UAS-Bnl:GFPΔFGF:* ATGAGCTGTACAAGACGGTGCCGCAGGAG | This paper | N/A |
| Forward primer for cloning all the constructs above: AATTCGAGCTCGGTACAGATCTATGCGAAGAAACCTGCG C | This paper | N/A |
| Primer for cloning *UAS-sBtl:Cherry*: AATTCGAGCTCGGTACCTCGAGATGGCAAAAGTGCCGAT CACG | This paper | N/A |
| Primer for cloning *UAS-sBtl:Cherry*: GCCGCCTTGCCCCTCGACAGGATGGGCGTGCAGCAG | This paper | N/A |
| Primer for cloning *UAS-sBtl:Cherry*: GTCGAGGGGCAAGGCGGCatggtgagcaagggcgag | This paper | N/A |
| Primer for cloning *UAS-sBtl:Cherry*: GCCAAGCTTGCATGCCTCTAGAttacttgtacagctcgtccatgcc | This paper | N/A |
| Forward primer for *bnl* RT-PCR: CAGGAGGACACTCACAATTGCCAG | This paper | N/A |
| Reverse primer for *bnl-RA* RT-PCR: GCTGCAGACACAGGAAATCG | This paper | N/A |
| Reverse primer for *bnl-PC* RT-PCR: GGGACAACAGTCCGAAATCG | This paper | N/A |
| Primer for cloning *UAS-Bnl*$^{PC}$*:GFP:* GCCAAGCTTGCATGCCATATATTCTAGATCATCGCCGGG GGGACAACAGTCCGAAATCGTAGTAGAGCGAATCGTCG | This paper | N/A |
| Recombinant DNA | | |
| pUAST-Bnl:GFP | [2] | N/A |
| pUAST-Bnl:HA | [2] | |
| pUAST-Bnl:HA$_1$GFP$_3$ (UAS-HA$_1$Bnl:GFP$_3$) | [2] | |
| pUAST-GFP-GPI | [7] | N/A |
| pUAST-cSpi:GFP | [8] | N/A |
| pUAST-Bnl:GFPΔC$_{40}$ (ΔC) & -Bnl:GFPΔC$_{168}$ | *UAS-Bnl:HA$_1$GFP$_3$* by deleting the last 40 (after Y$_{730}$ of Bnl) and 168 (after K$_{602}$ of Bnl) amino acid regions, respectively, prior to a stop codon. | N/A |
| pUAST-Bnl:GFPΔC-TM & -Bnl:GFPΔC-TM$_{168}$ | a 31 amino acid long transmembrane domain of the mammalian CD8a protein fused to the C-terminus of *UAS-Bnl:GFPΔC* and *UAS-Bnl:GFPΔC$_{168}$*, respectively. | N/A |

| | | |
|---|---|---|
| pUAST-Bnl:$GFP_3Cherry_c$ | *UAS-Bnl:$HA_1GFP_3$* with a C-terminal mCherry tag with a linker (VEGQGG) placed in between. | N/A |
| pUAST-$Bnl_{PC}$:GFP | The PC-specific C-terminal 24 bp sequence (7 amino acids+stop) was added to the C-terminus of the 1-2259 bp region of *bnl-PA* CDS using PCR | N/A |
| pUAST-Bnl:GFP-$\omega^m$ | *UAS-Bnl:$HA_1GFP_3$* with mutated $\omega$, $\omega+1$, and $\omega+2$ sites (S/$P^{741}$G/$P^{742}$A/$P^{743}$) | N/A |
| pUAST-bGFP-GPI | secGFP [1] (superfolder GFP with N-terminal Bnl signal peptide) added with the last 53 amino acids of Bnl (from $T_{718}$) at the C-terminus. | N/A |
| pUAST-sBtl:Cherry | mCherry sequence was added in-frame after $P^{607}$ of Btl, replacing the TM and intracellular portions. | N/A |
| pUAST-BtlDN:Cherry and pLot-BtlDN:Cherry | mCherry sequence was added in-frame after $L_{625}$ of Btl, replacing the intracellular C-terminal portions. | N/A |
| pUAST-Bnl:$GFP^{\Delta FGF}$ | Conserved FGF domain of Bnl was replaced with a sfGFP sequence | N//A |
| Software and Algorithms | | |
| Fiji | ImageJ | https://fiji.sc |
| Prism 8.0 | GraphPad | https://www.graphpad.com/ |
| Adobe Photoshop | Adobe | https://www.adobe.com |
| Adobe Illustrator | Adobe | https://www.adobe.com |
| Microsoft Excel | Microsoft | https://www.office.com |
| SnapGene | SnapGene | https://www.snapgene.com |
| MacVector | MacVector | https://macvector.com |
| PredGPI predictor | | http://gpcr.biocomp.unibo.it/predgpi/pred.htm |
| VassarStats | | vassarstats.net |
| R x64 3.3.1 | R | r-project.org |
| Imaris 9.5.0 | Imaris | https://imaris.oxinst.com |

#### Supplementary Notes

##### A. Expression analyses of *bnl* splice variants using RT-PCR

The *bnl* gene has two different splice variants encoding proteins with different C-terminal hydrophobic sequences. To check the expression of different *bnl* isoforms, total RNA was extracted from 20 $w^{1118}$ larval wing discs and RT-PCR was performed on the total RNA. For both isoforms, we used a common forward primer that binds to exon 3 (5'-CAGGAGGACACTCACAATTGCCAG-3'). However, the reverse primers were either PA- (5'-GCTGCAGACACAGGAAATCG-3') and PC- (5'-GGGACAACAGTCCGAAATCG-3') -specific. The reverse primer for each isoform was designed to span the junction between exon 3 and the isoform-specific last exon as illustrated below. The expected amplicon size was 697bp. As a negative control, RT-PCR was carried out on the genomic DNA (gDNA) template obtained from $w^{1118}$ flies. As a positive control we performed RT-PCR for constitutive *rp49* gene. Unsurprisingly, RT-PCR results showed strong amplification of *bnl-PA,* exclusively from the RNA template. Although we detected strong *bnl-PC* amplification from the RNA template, we also detected a low level amplification of the same sized RT-PCR product from gDNA. These results confirmed *bnl-PA* expression in the wing disc. These results, although not conclusive, also suggested that the wing disc expresses *bnl-PC*. Moreover, a Bnl antibody, which detects both isoforms, showed that the native $Bnl^{ex}$ is asymmetrically localized on the wing disc producing cell surface and the $Bnl^{ex}$ was reduced with the PIPLC treatment. Secondly, S2 cells expressing a chimeric $Bnl^{PC}$:GFP construct showed the PIPLC-sensitive surface distribution of the protein. Based on these results, we suggest that irrespective of the tissue-specific expression levels, Bnl isoforms are GPI-anchored on the cell surface. Consistent RT-PCR results were obtained from three independent experiments, confirming the expression profile.

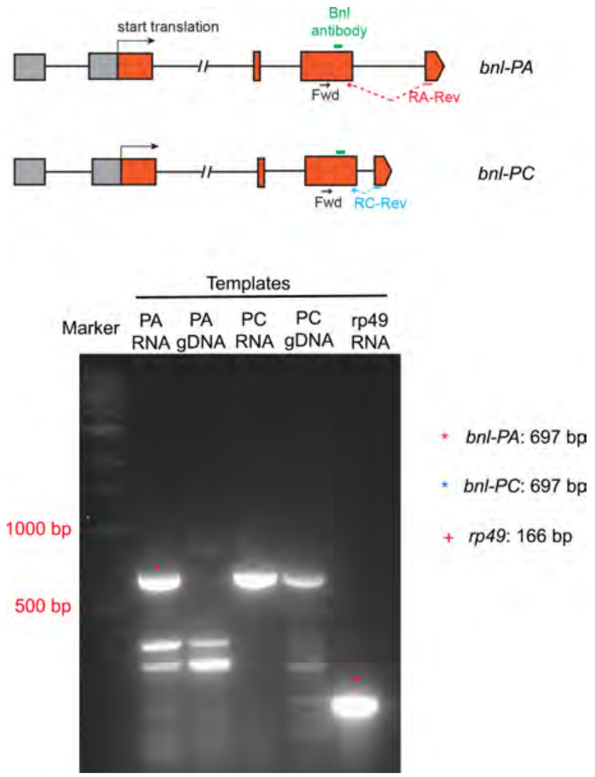

#### B. Bioinformatic analyses of hydropathy and secondary topology of various BnI constructs

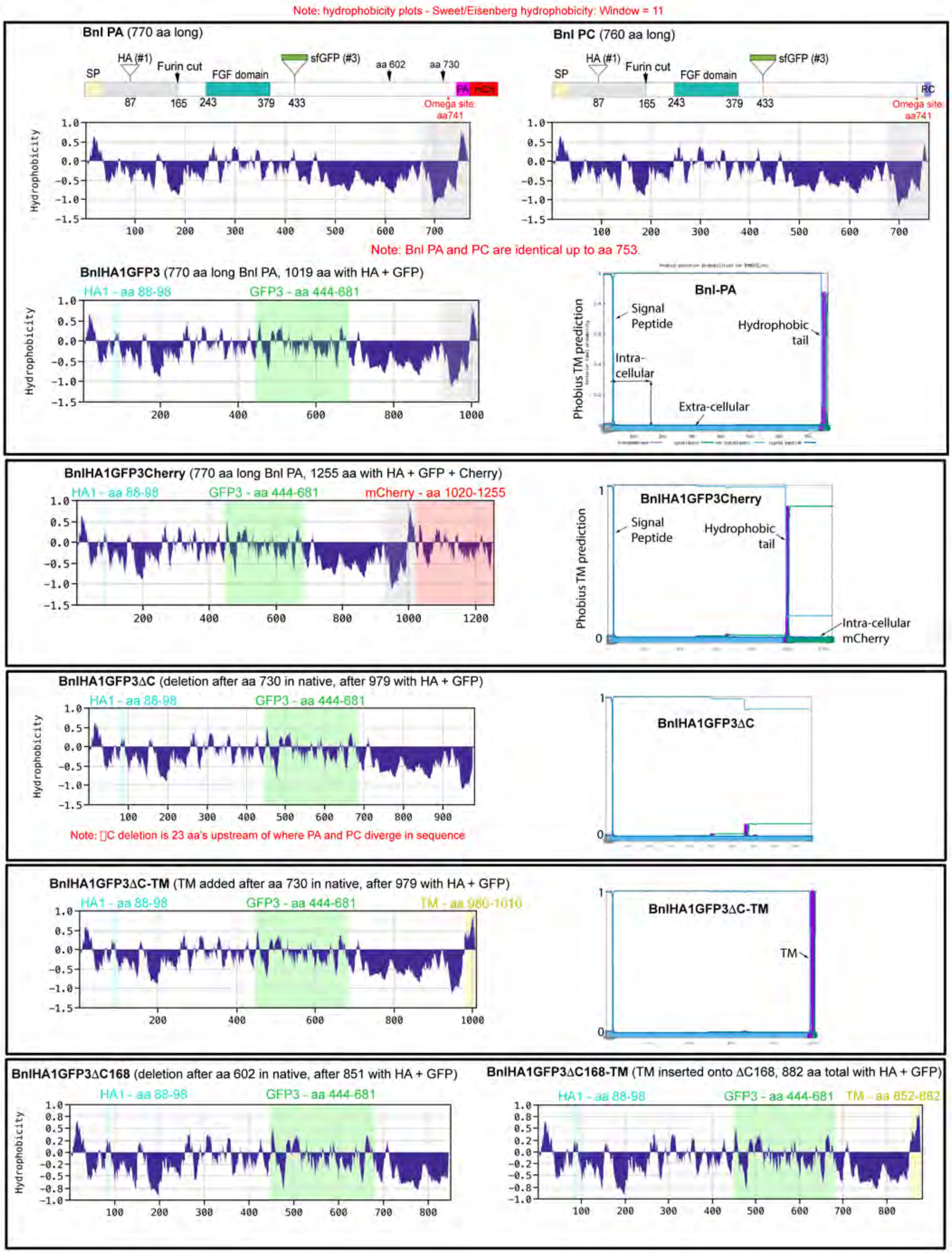

#### C. Comparison of *bnl-GAL4*-driven expression levels of transgenic constructs

| Transgenic Constructs | Fly line | Chromosomal Insertion | Expression levels * | externalized by source? | ASP uptake? |
|---|---|---|---|---|---|
| UAS-Bnl:GFPΔC40 | 2_2 | 3 | +++ | Yes | Yes |
| | 3_1 | 3 | ND | n/a | n/a |
| | 3_2 | 2 | ++ | Yes | Yes |
| UAS-Bnl:GFPΔC40-TM | 1_2 | 3 | +++ | Yes | Yes (R) |
| | 2_2 | 2 | +++ | Yes | Yes (R) |
| | 3_1 | 3 | ++ | Yes | Yes (R) |
| | 3_2 | 2 | ++ | Yes | Yes (R) |
| | 4_1 | 2 | ++ | Yes | Yes (R) |
| | 6_1 | 3 | ND | n/a | n/a |
| | 7_1 | 3 | + | Yes | Yes (R) |
| | 9_2 | 3 | +++ | Yes | Yes (R) |
| UAS-Bnl:GFPΔC168 | 1_2 | 2 | ++ | Yes | Yes |
| | 4_1 | 3 | +++ | Yes | Yes |
| UAS-Bnl:GFPΔC168-TM | 1_1 | 3 | ++ | Yes | Yes (R) |
| | 2_1 | 3 | +++ | Yes | Yes (R) |
| UAS-Bnl:GFP-$\omega^m$ | 1_1 | 3 | +++ | L | L |
| | 4_1 | 2 | +++ | L | L |

* Expression levels were verified by *bnl-Gal4* driven expression of the *UAS* constructs in the wing disc *bnl* source (see Methods). Bnl:GFP lines used in this study was published earlier in Du et al. and Sohr et al.[1,2]. ND: Not detected. Red: Lines with comparable expression levels used in this study. R: Restricted range. L: Very low level.

+++ >++>+ : High>medium>low levels of expression relative to each other.

#### An example of comparison of levels of expression of Bnl:GFP variants under *bnl-Gal4* in the wing disc

| | Bnl:GFP | Bnl:GFPΔC40 (2_2) | Bnl:GFPΔC40-TM (2_2) |
|---|---|---|---|
| Mean GFP intensity in the *bnl*-source (derived from extended Z-stack of 50 μm tissue) (7 wing discs from 7 animals used) | 1137.40 | 867.5 | 1495.3 |
| | 1142.2 | 1106.3 | 899.7 |
| | 1870.1 | 2400.1 | 620.6 |
| | 747.7 | 909.8 | 1372.5 |
| | 1190.05 | 782.9 | 610.1 |
| | 972.8 | 985.5 | 1868.6 |
| | 527.6 | 484.3 | 1126.298 |
| **Average** | **1083.98*** | **1076.63*** | **1141.87*** |

*, Comparable levels of expression of different constructs used in this study.

**D. Examples of gating strategy for FACS analyses**

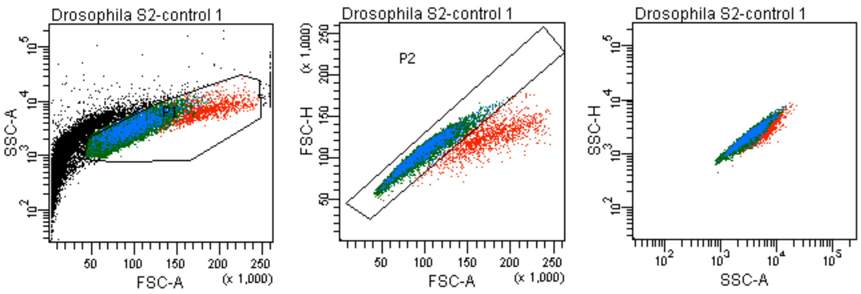

```
library(ggplot2)

filename="Control Source"

file_t= "data\\"
file<-paste(file_t,filename,".csv",sep="")

#data<-read.csv("FinalData\\Control Source.csv",head=TRUE,sep=",")
data<-read.csv(file,head=TRUE,sep=",")

cols<-c("#619cff","#f8766d","#00ba38")
#cols<-c("red","blue","green")

p<-ggplot(data, aes(x=data$Range,y=data$Counts,fill=data$Length)) +
 #theme_bw()+
 #theme_minimal() +
 geom_bar(width = 30, colour="black", stat="identity") +
 #geom_hline(yintercept = 2.5) +
 #geom_vline(xintercept = c(0,90,180,270)) +
 scale_fill_manual(values = cols) +
 #scale_y_discrete(drop = FALSE) +
 theme(legend.box.just = "top",legend.position = "bottom") +
 theme(panel.grid.major = element_line(colour = "gray"),
        panel.grid.minor = element_line(colour = "blue"),
        panel.background = element_blank(),
        axis.line = element_line(colour = "black"))+
 labs(title = filename,
        fill = "Length range(um)",
        y = "Cytoneme number", limits = c(0, 100),colour =
"Cylinders") +
 coord_polar(theta = "x", start=pi/2, direction=-1) +
 scale_x_discrete("",  limits = c(0,90,180,270), labels =
c(0,90,180,270)) +
 scale_y_continuous(limits=c(0,5), breaks= c(1:5))


p
```
